# Regulation of mitochondrial dynamics by Aurora A kinase

**DOI:** 10.1101/236612

**Authors:** Rhys Grant, Ahmed Abdelbaki, Alessia Bertoldi, Maria P Gavilan, Jörg Mansfeld, David M Glover, Catherine Lindon

## Abstract

Aurora A kinase (AURKA) is a major regulator of mitosis and an important driver of cancer progression. The roles of AURKA outside of mitosis, and how these might contribute to cancer progression, are not well understood. Here we show that a fraction of cytoplasmic AURKA is associated with mitochondria, co-fractionating in cell extracts and interacting with mitochondrial proteins by reciprocal co-immunoprecipitation. We have also found that the dynamics of the mitochondrial network are sensitive to AURKA inhibition, depletion or overexpression. This can account for the different mitochondrial morphologies observed in RPE1 and U2OS cell lines, which show very different levels of expression of AURKA. We identify the mitochondrial fraction of AURKA as influencing mitochondrial morphology, since an N-terminally truncated version of the kinase that does not localize to mitochondria does not affect the mitochondrial network. We identify a cryptic mitochondrial targeting sequence in the AURKA N-terminus and discuss how alternative conformations of the protein may influence its cytoplasmic fate.

## Introduction

AURKA was discovered as a mitotic kinase with a key role in bipolar spindle formation (Glover et al., 1995). Its prominent localization to the microtubule spindle in mitosis is mediated through interaction of its kinase domain with the microtubule associated protein (MAP) TPX2 (Kufer et al., 2002), an interaction that also activates the kinase activity of AURKA through stabilization of the active conformation of the T-loop (Dodson and Bayliss, 2012). Further studies have revealed the existence of alternative pathways to activation of AURKA (Joukov et al., 2010; Plotnikova et al., 2010; Reboutier et al., 2012) and an extensive array of cellular functions regulated by AURKA, in interphase as well as mitosis (reviewed in (Nikonova et al., 2013)). One such novel function that has been described for AURKA is the promotion of mitochondrial fission in preparation for mitosis (Kashatus et al., 2011).

Mitochondria form a highly dynamic network of interconnected tubules that undergo constant cycles of fission and fusion. These cycles are an essential feature of cellular homeostasis, thought to maintain a healthy mitochondrial population by allowing segregation of damaged mitochondria into the mitophagy pathway (Twig et al., 2008; Youle and van der Bliek, 2012). It is also proposed that fission/fusion cycles regulate the metabolic output of the cell, since a more interconnected network provides more efficient oxidative metabolism, and that mitochondrial morphology responds to metabolic cues. Mitochondrial morphology depends on the activity of GTPases dynamin-related protein 1 (Drp1) for fission, and mitofusins (MFN1, MFN2) for fusion, of the outer mitochondrial membrane (OMM) (Scott and Youle, 2010; Smirnova et al., 2001).

Mitochondrial fission also occurs prior to cell division in mammalian cells. Fragmentation of organelle networks (mitochondria, Golgi, ER) is a common feature of cell division, thought to improve the probability that organelles segregate equally between daughter cells (Jongsma et al., 2015; Taguchi et al., 2007) and to enable regulated segregation of aged mitochondria during stem cell-like divisions (Katajisto et al., 2015). Mitochondrial fragmentation in mitosis may also be required for efficient microtubule-dependent transport of mitochondria towards the cell periphery and away from the cell division plane (Kanfer et al., 2015). Indeed, blocking mitochondrial fission or transport gives rise to cytokinesis failure and aneuploidy (Mitra et al., 2012; Qian et al., 2012; Smirnova et al., 2001), an effect that can be rescued by disrupting mitochondrial fusion (Rohn et al., 2014). Previous studies have described a pathway leading to increased mitochondrial fission at mitotic entry via Cdk1-dependent activation of Drp1 at the OMM (Taguchi et al., 2007). The mitotic kinase AURKA has been proposed to regulate these events through phosphorylation of RalA to promote RalA-dependent OMM recruitment of both Drp1 and RalBP1-cyclinB-Cdk1-complex (Kashatus et al., 2011).

Fragmented mitochondrial networks are a characteristic of cancer cells thought to contribute to the metabolic changes that accompany tumorigenesis, and have been shown to contribute to tumour progression and metastasis (Regan et al., 2013; Ward and Thompson, 2012; Zhao et al., 2013). Overexpressed AURKA (located on the 20q amplicon) is also strongly associated with cancer (Bischoff et al., 1998; Zhou et al., 1998). Given the large number of mitochondrial hits we identified in a search for AURKA interactors, we decided to investigate further the role of AURKA in influencing the fragmentation state of the mitochondrial network in human cell lines. We report here that the mitochondrial network is sensitive to AURKA activity at all phases of the cell cycle, and that this sensitivity contributes to divergent mitochondrial morphology between two cell lines (one transformed and the other non-transformed) expressing different levels of AURKA. Furthermore we identify an interphase subpopulation of AURKA associated with mitochondrial fragments, both by immunofluorescence and fractionation assays. This association depends on the AURKA N-terminal region, which contains a cryptic mitochondrial targeting sequence.

## Materials and Methods

### Plasmids

pVenus-N1-AURKA has been previously described (Min et al., 2013). Deletion mutants were made using standard cloning and site directed mutagenesis techniques and verified by sequencing. Full cloning details are available on request. pcDNA5-FRT/TO-AURKA-Venus was made from pcDNA5™/FRT/TO (Thermo Fisher Scientific) with HygromycinR sequences replaced by NeomycinR.

### Cell culture and treatments

hTERT-RPE-1 (RPE-1) or RPE-1 FRT/TO cells (Mansfeld et al., 2011) cells were cultured in 50:50 mix of Ham’s F12:DMEM and U2OS cells in high glucose DMEM (Thermo Fisher Scientific). All cell culture media were supplemented with FBS (10%), Penicillin-Streptomycin and amphotericin B. RPE-1-AURKA-Venus (Flp-In) lines were derived as polyclonal populations by pooling cells transfected with pcDNA5-FRT/TO-AURKA-Venus and Flp-recombinase (pOG44) after 12 days selection in 500 ug mL^−1^ geneticin. Expression of AURKA-Venus is achieved by supplementing cell culture medium with 1 ug mL^−1^ tetracycline hydrochloride (tet, Calbiochem, San Diego, CA, USA). U2OS-AURKA-Venus cells were obtained by clonal selection with 170 mg ml^−1^ hygromycin B (Calbiochem) after transfection of U2OS tet-OFF cells with pTRE-AURKA-Venus:pCMVhygro in a ratio of 10:1. Expression of AURKA-Venus in these cells is achieved with extensive washing in PBS and switch to tet-free cell culture medium. To generate the Aurora A-mVenus knock in RPE-1 FRT/ TO cells were infected with recombinant adeno-associated virus particles harboring mVenus cDNA flanked by ~1,500 bp homologous to the *AURKA* locus as previously described (Collin et al., 2013). To identify positive integrands cells were treated with 10 mM DMA 12 hours before single cell sorting by flow cytometry. mVenus positive cells were verified by fluorescence microscopy and immunoblot analysis (Supplemental figure S2). D.mel-2 cells were cultured at 25°C in Express Five^®^ serum free medium supplemented with antibiotics and 2mM L-glutamine.

siRNA duplex targeting human AURKA, and control siRNA duplex against GL2, have been previously described (Floyd et al., 2008)(Sigma-Aldrich).

Transfections were carried out using Invitrogen Neon^®^ system to electroporate cells with siRNA or plasmids, according to the manufacturers’ instructions.

For mitochondrial imaging in living cells, cells were incubated in 100 nM MitoTracker™ Red CMXRos (Thermo Fisher Scientific) or Mito-ID^®^ green (Enzo) diluted in filming medium (see below) for 15 min. MitoTracker™/Mito-ID^®^ were replaced with fresh filming medium prior to timelapse.

Cells were treated with 100 nM MLN8237 during time-lapse experiments, by addition of drug in a volume not less than 1/100 of the existing dish volume and gentle mixing of medium on the dish.

### Cell extracts, fractionation and immunoblotting

Whole cell extracts were prepared by scraping cells directly into 2X SDS Sample Buffer with 10 mM DTT. Samples were syringed to shear DNA or sonicated and boiled at 95 °C for 5 minutes prior to SDS-PAGE on 4-12% pre-cast gradient gels (Invitrogen). Transfer onto Immobilon-P or Immobilon-FL membranes was carried out using the XCell IITM Blot Module according to the manufacturer’s instructions. Membranes were blocked in PBS/ 0.1% Tween-20/ 5% dried milk and processed for immunoblotting with primary antibodies indicated in figures. Secondary antibodies used were HRP-conjugated, or IRDye^®^ 680RD- or 800CW-conjugated for quantitative fluorescence measurements on an Odyssey^®^ Fc Dual-Mode Imaging System (LI-COR Biosciences). For cell fractionation experiments, cells were harvested and treated at 4 °C with 40 μg/mL digitonin in 10 mM Tris-HCl pH7.4, 100 mM NaCl, 25 mM MgCl_2_, 1 mM Na_3_VO_4_, 1 mM NaF, EDTA-free protease inhibitor cocktail (Roche). Cells were disrupted by 10 passages through a 25 gauge needle (whole cell extract). Cell nuclei were pelleted by centrifugation at 3000 rpm/ 10 min in a microfuge at 4 °C. The resulting supernatant was recentrifuged at 13000 rpm/ 15 min to yield mitochondrial (pellet) and cytoplasmic (supernatant) fractions.

Quantitative immunoblotting was carried out using IRDye^®^ 680RD and 800CW fluorescent secondary antibodies, scanned on an Odyssey^®^ Imaging System (LI-COR Biosciences).

### Immunofluorescence analysis

Cells were grown on 13 mm glass coverslips, synchronized and fixed using either 100% MeOH at –20 °C or 4% PFA treatment at ambient temperature. Mitochondria were detected using anti-TOMM20 (Santa Cruz), AURKA using anti-AURKA (rabbit antibody Abcam ab1287 or mouse antibody BD Transduction Laboratories 610939) and AURKA-Venus using anti-GFP (Abcam ab290). Secondary antibodies used were Alexa®Fluor 488-anti-rabbit and Alexa®Fluor 568-anti-mouse (Thermo Fisher Scientific). Cells were stained according to standard protocols in PBS buffer containing 0.1% Triton X100 and 3% BSA and mounted in ProLong^®^ Gold antifade (Thermo Fisher Scientific). Images were captured using an Axiovert 200M fluorescence microscope (Carl Zeiss Inc.) with a 100X NA 1.4 oil objective and Coolsnap HQ2 camera (Photometrics) controlled by MetaMorph^®^ software (MDS ABalytical Technologies). Images were deconvolved using 10 iterations of AUTOQuant X2 (Media Cybernetics) blind deconvolution algorithm and presented as maximum intensity projections of 10 x 0.1 um stacks using ImageJ. Intensity level, contrast and brightness of images were adjusted using Adobe Photoshop where indicated.

### Live cell microscopy

Cells were seeded onto 8-well plastic-bottom slides (Ibidi GmbH, Martinsried, Germany) at density of 8 x 10^4^ cm^−2^. Imaging medium was Leibowitz L-15 (Thermo Fisher Scientific) supplemented with FBS and antibiotics as described above. Expression of AURKA-Venus was induced 24 hours prior to imaging, which was carried out on an Olympus CellR widefield imaging platform comprised of Olympus IX81 motorized inverted microscope, Orca CCD camera (Hamamatsu Photonics, Japan), motorized stage (Prior Scientific, Cambridge, UK) and 37°C incubation chamber (Solent Scientific, Segensworth, UK) fitted with appropriate filter sets and 60X NA 1.42 oil objective. Images were acquired using Olympus CellR software, as 1 um stacks and exported as 12 bit TIFF stacks for display as maximum intensity projections in ImageJ.

### Mitochondrial morphology analysis

MitoTracker^™^ images were analysed using both manual analysis with ImageJ software and automated analysis using MicroP software (Peng et al., 2011). For manual analysis, the lengths of 30 mitochondria per cell were measured, starting at the 12 o’clock position and moving clockwise around the nucleus. Results from manual analysis are presented in figures, using kernal density estimations to derive probability density plots in R. In each experiment, mean mitochondrial tubular lengths have been calculated using MicroP and corroborated as statistically significant, as indicated in figure legends.

## Results and Discussion

In using proteomic approaches to identify co-purifying proteins in AURKA-GFP pulldowns from human U2OS cells, we identified large numbers of mitochondrial proteins (Supplementary Figure S1A). We confirmed these potential interactions by reciprocal pulldowns of several GFP-tagged mitochondrial markers, including the OMM translocase complex components TOMM20 and TOMM70 and the inner mitochondrial membrane (IMM) component Prohibitin (PHB), which identified endogenous AURKA as a partner protein in cell extracts (Figure S1B). Repeated experiments suggested that the quantity of AURKA in pulldowns of mitochondrial proteins was at least 10-fold less than that found in pulldowns of TPX2, a major interactor of AURKA in mitotic cells (Kufer et al., 2002).

Given the previous study suggesting that mitochondrial fission was controlled by a cytoplasmic pool of AURKA (Kashatus et al., 2011), we investigated the relevance of our own finding - of the association of AURKA with mitochondrial proteins - to mitochondrial morphology. First we tested whether depletion of endogenous AURKA, using siRNA-mediated knockdown (AurAi), would affect mitochondrial morphology in the immortalized Retinal Pigment Epithelial line hTERT-RPE-1 (RPE-1). We found that AurAi resulted in markedly reduced fragmentation of the mitochondrial network, observable as a more interconnected network and quantifiable as a change in length distribution of individual mitochondrial fragments (Figure 1A). To confirm the specificity of this effect, we treated RPE-1 cells, expressing a mCherry-PCNA marker to distinguish cell cycle phases, with the small molecule specific inhibitor of AURKA, MLN8237. We found that the length of mitochondrial fragments varied with cell cycle phase, as expected (Mitra et al., 2009; Taguchi et al., 2007), being shortest in G1 phase (Figure 1B), but AURKA inhibition led to elongation of mitochondria in all cell cycle phases (Figure 1C). Therefore we conclude that AURKA can regulate mitochondrial organization throughout the cell cycle. To determine whether this response of mitochondria to AURKA is a conserved role of the kinase, we treated cultured *D. melanogaster* DMEL cells with MLN8237 (Figure 1D). The increased in mitochondrial connections and length following this treatment suggest that this role for AURKA is conserved in metozoans.

**Figure 1.**
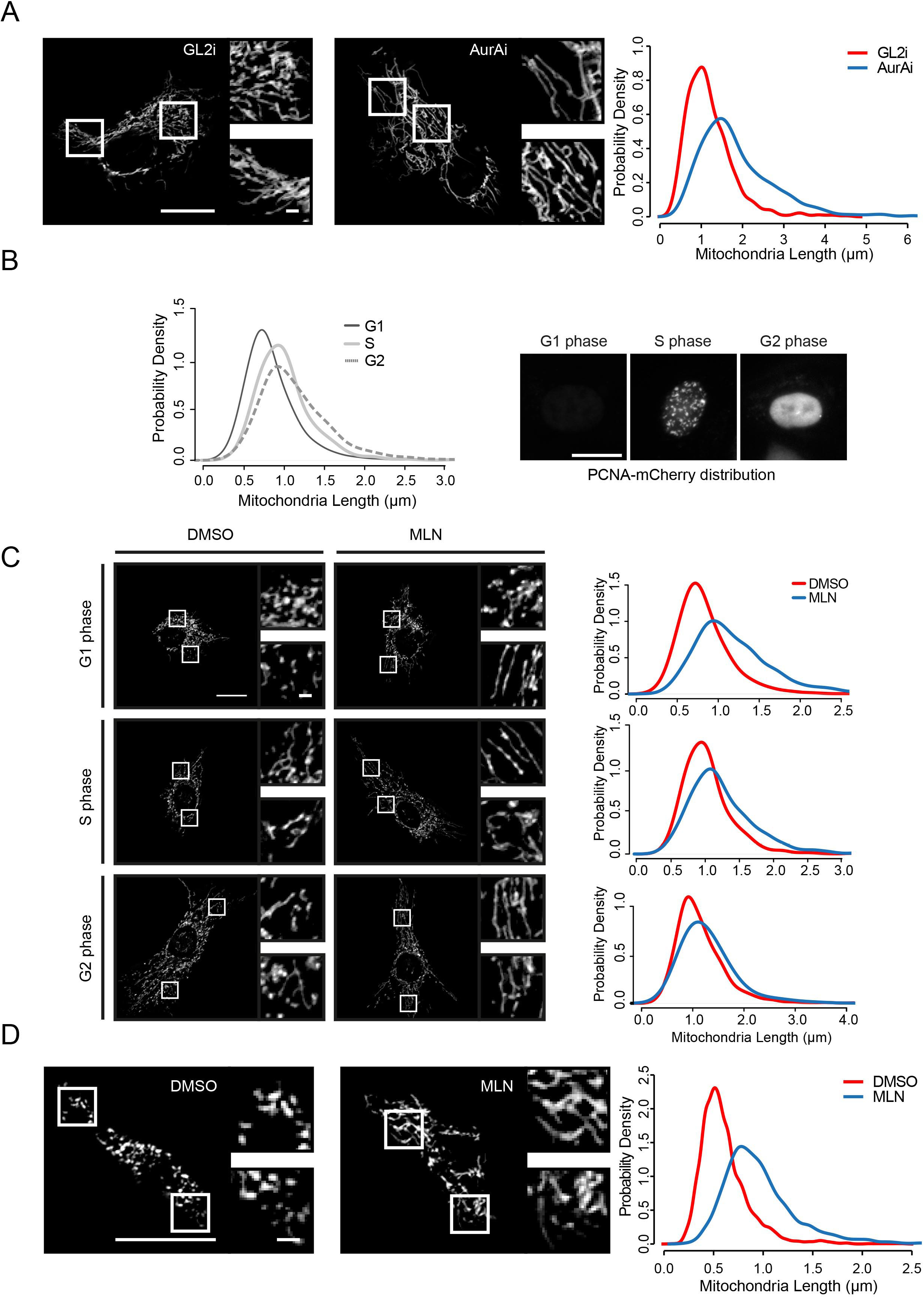
AURKA is a constitutive regulator of the mitochondrial network. **A** RPE-1 cells were treated with control (GL2i) or AURKA siRNA (AurAi) for 48 h and mitochondria imaged in live cells using MitoTracker^TM^. Areas of cytoplasm marked by white squares are shown enlarged twofold in panels to the side of each image. Mitochondria were manually analysed for tubular fragment size as described in Material and Methods and results shown as probability density plots. **B,C** RPE-1-PCNA-mCherry cells were stained with Mito-ID^®^ green and imaged after treatment with 100 nM MLN8237 (MLN) or vehicle control for 3 hours. Cells were assigned to G1, S or G2 phase according to PCNA marker (Materials and Methods) and analysed for tubular mitochondrial lengths as in **A. B** Probability density plots of mitochondrial length according to cell cycle phase. **C** Probability density plots showing increased mitochondrial fragment length in all phases of the cell cycle after MLN treatment. **D** Drosophila S2 cells were treated with MLN for 3 hours and processed as in **A.** For all parts **A-D**: Scale bars, 10 μm in main panels, 1μm on magnification panels. n ≥ 20 cells, *p* < 0.001 (Mann-Whitney test), 3 repeats.

We noticed that two cancer cell lines commonly used in our lab (U2OS and HeLa) both had mitochondrial networks that always appeared in a more fragmented state than in nontransformed RPE-1 cells (Figure 2A, data not shown). Given our finding that AURKA influences mitochondrial morphology, and the well-documented overexpression of AURKA in cancer cells, we tested whether AURKA expression levels might contribute to differences in mitochondrial morphology observed between RPE-1 and U2OS cells. We determined relative expression levels in extracts prepared from known numbers of cells in asynchronous populations, normalized against a panel of cellular proteins (Figure 2B). This revealed that U2OS cells contain two-fold higher levels of AURKA protein normalised against cell number, and 4-fold to 6-fold more when normalised against the levels of tubulin, actin or ATP5A1. Therefore, higher AURKA levels correlate with a more fragmented mitochondrial network. Next we tested whether manipulating AURKA levels would reproduce observed patterns of mitochondrial organization. We used partial siRNA-mediated knockdown to achieve 6-fold reduction of AURKA levels in U2OS cells, and found a corresponding lengthening of mitochondrial fragments under these conditions (Figure 2C,E). Connectivity of the network was also greater when AURKA levels were reduced. Conversely, we found that tetracycline-induced overexpression of an AURKA-Venus transgene in RPE-1 cells resulted in increased fragmentation of the mitochondrial network (Figure 2D,E). Therefore we conclude that the different fragmentation state of the mitochondrial network in different cell lines is a direct reflection of the level of AURKA expression.

**Figure 2.**
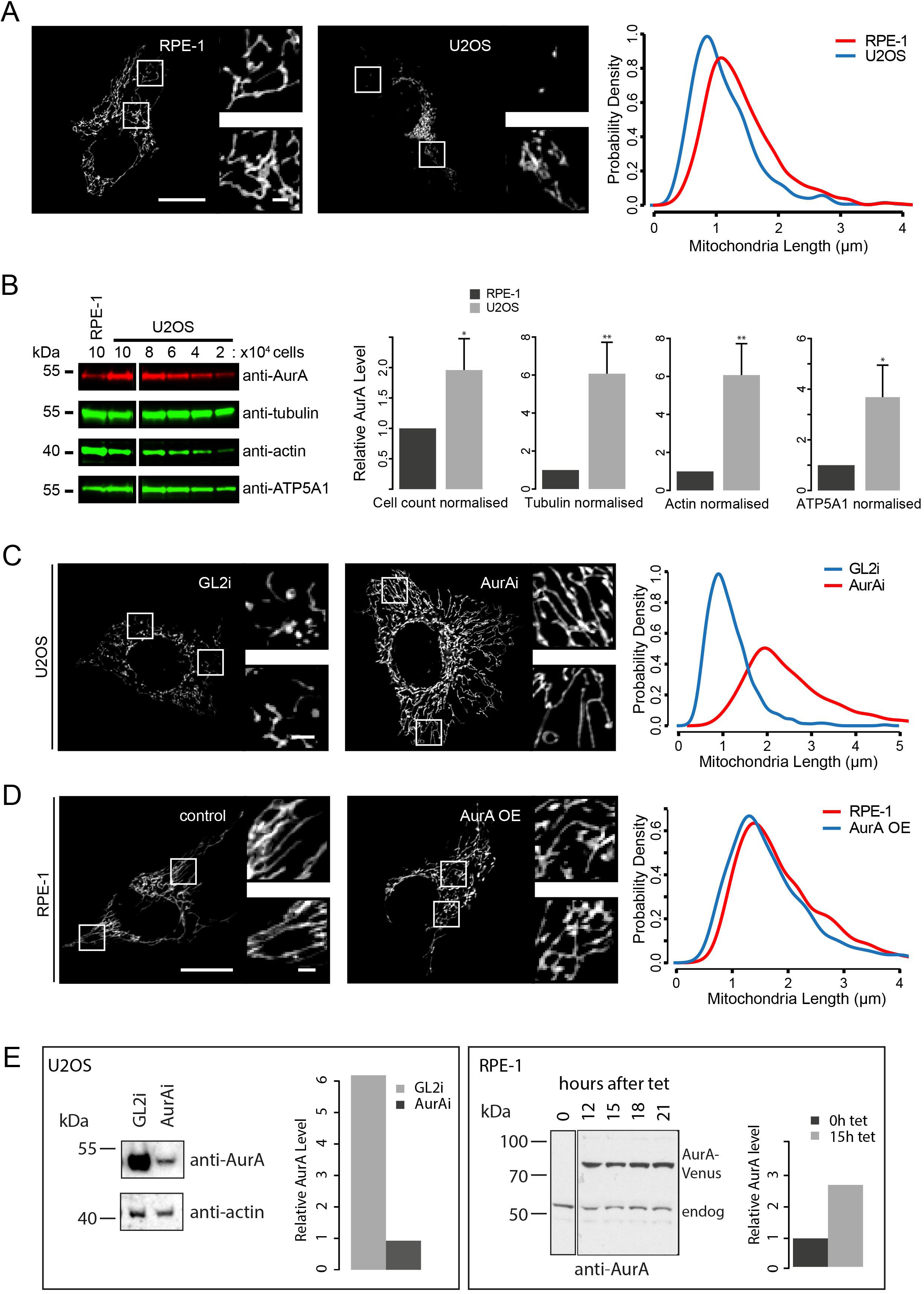
AURKA levels influence mitochondrial morphology in RPE and U2OS cells. **A** RPE-1 and U2OS were treated with MitoTracker^™^ and imaged under identical conditions. Mitochondrial lengths were measured and plotted as in Figure 1. n = 9 – 18 cells, *p* < 0.001 (Mann-Whitney U-test), 2 repeats. **B** Equal numbers of U2OS and RPE-1 cells were harvested for cell extracts and calculated quantities loaded onto gels to be examined by quantitative immunoblot. AURKA levels were quantified and normalized against number of cells, or against different loading markers. **C-E** U2OS cells were treated with control (GL2i) or AURKA siRNA for 48h (**C**), whilst RPE-1-AURKA-Venus cells were induced for AURKA-Venus expression (AURKAoe) or not (control) with addition of tet for 18 hours (**D**). Cells were either processed for quantitative immunoblotting of AURKA levels (**E**) or stained with MitoTracker^™^ for mitochondrial tubular length measurements as in **A** (**C, D**). OE, overexpression; tet, tetracycline.

Given the prevalence of mitochondrial hits in our AURKA interactome, and the functional relationship between AURKA and mitochondrial organization, we hypothesized that part of the cytoplasmic pool of AURKA might be associated with the mitochondrial network. In support of this, we found that both endogenous and exogenous AURKA co-purify with mitochondria in extracts fractionated by centrifugation (Figure 3A). We also examined fixed cells for colocalization of endogenous AURKA with a mitochondrial marker, TOMM20. Immunostaining with two different antibodies against AURKA revealed punctate cytoplasmic staining in which the AURKA “dots” were frequently apposed to mitochondrial fragments, often close to points of mitochondrial constriction (Figure 3B). We then sought to confirm that these dots were AURKA-specific using siRNA to deplete endogenous AURKA. This led to extensive reduction of the punctate staining associated with mitochondria in accord with the down-regulation of AURKA by AurAi treatment (Supplementary Figure S1C,D). To confirm this we examined the cytoplasmic localization of exogenous AURKA-Venus in living cells (that is, without fixation). When AURKA-Venus was expressed from a tet-ON promoter, levels of cytoplasmic AURKA were too high to distinguish detailed staining patterns from the high cytoplasmic background (Fig. 1C). To circumvent this issue we targeted one allele of endogenous AURKA at the C terminus with mVenus taking advantage of rAAV-mediated homologous recombination (Supplemental Figure S2A). Endogenous mVenus-tagged AURKA was expressed to the same level as untagged endogenous AURKA and was comparably enriched cells arrested in mitosis by dimethylenastron (DMA) (Fig. S2B,C). Notably, AURKA-mVenus localized to the centrosomes in G2 phase, to the mitotic spindle, midzone and midbody in mitosis and was degraded from the onset of anaphase (Fig. S2D). Together this suggesting that AURKA-mVenus faithfully recapitulates the known cell cycle behaviour of untagged AURKA. When we examined endogenous AURKA-mVenus, which was expressed at approximately 10-fold lower levels than from the tet-ON promoter, we could detect a similar pattern of colocalization of cytoplasmic dots of Venus fluorescence with constrictions in mitochondria in the living cells as we had seen when staining fixed cells for endogenous AURKA (Figure 3C,D). Therefore we conclude that a small fraction of cytoplasmic AURKA resides at mitochondria.

**Figure 3.**
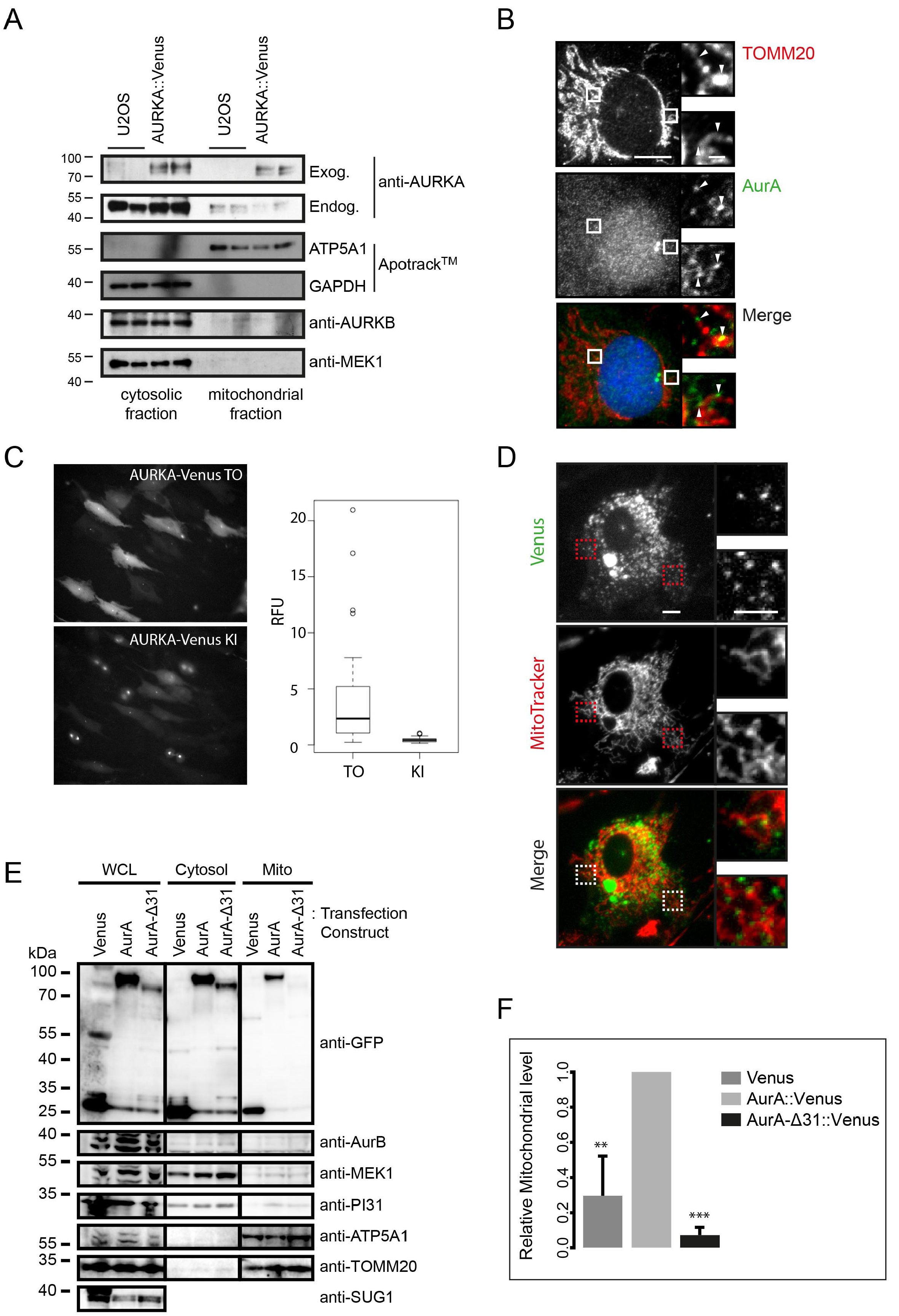
Mitochondrial fraction of AURKA depends on N-terminal sequences. **A** U2OS and U2OS-AURKA-Venus cells were fractionated by serial centrifugation and probed for the presence of exogenous and endogenous AURKA, and of specific markers for cytosolic (GAPDH, MEK1) or mitochondrial (ATP5A1) fractions, by immunoblot. **B** Colocalization of endogenous AURKA (green) and mitochondrial marker TOMM20 (red) examined by IF in RPE-1 cells. **C** Comparison of AURKA-Venus levels in endogenously tagged RPE-1 Knock-In (KI) cells and RPE-1-AURKA-Venus Tet-On (TO) cells, by quantitative measurements of Venus fluorescence in living cells. Box and whisker plots show mean fluorescence values in cells expressing AURKA-Venus, n > 45, *p* < 0.0001 (Student’s t-test). **D** Live cell imaging of KI cells stained with MitoTracker™ to examine colocalization with endogenously tagged AURKA. **E,F** U2OS cells transiently transfected with AURKA-Venus (AurA), N-terminally truncated AURKA (AurA-Δ31) or Venus alone were fractionated and probed by immunoblot for the presence of Venus (anti-GFP) or with various markers for cytosolic (MEK1, PI31) and mitochondrial (ATP5A1, TOMM20) fractions. SUG1 was used to control for Whole cell lysate (WCL). Venus levels in the mitochondrial fraction were quantitated and normalized against the level in WCL in 3 separate experiments. **, *p* < 0.01; ***, *p* < 0.001 (Student’s t-test).

The interaction we identified between TOM complex components and AURKA raised the possibility that AURKA might be targeted to mitochondria through a mitochondrial targeting sequence (MTS) (Horwich et al., 1985; Hurt et al., 1984). These are characterized by an amphipathic helix at the N-terminus. The known structure of AURKA (Bayliss et al., 2003) excludes the N-terminal region, which is generally considered to be unstructured. We tested if AURKA lacking its N-terminal 31 amino acids (AURKAΔ31) would still localize in the mitochondrial fraction in cell fractionation experiments and we found that it did not (Figure 3E,F). Moreover, AURKAΔ31 overexpression did not cause fragmentation of mitochondria (Supplementary Figure S2A). We conclude that the N-terminal region of AURKA contains sequences necessary for its localization with mitochondria and that this localisation is required for mitochondria to fragment in response to the kinase’s activity. To test this idea, we fused a recently described MTS, the N-terminal peptide (1-10) of survivin (BIRC5), (Dunajova et al., 2016) onto the N-terminus of AURKAΔ31. As predicted, expression of this gene fusion led to fragmentation of mitochondria (Supplementary Figure S2C).

These findings led us to ask whether the natural N-terminal domain of AURKA had any of the characteristics of mitochondrial targeting sequences. We found that AURKA’s N-terminal region is indeed polybasic and has the periodic hydrophobicity consistent with that of an amphipathic helix (Figure 4A). This predicted amphipathic helix is a phylogenetically conserved secondary feature, even where the primary amino acid (AA) sequence diverges (Supplementary Figure S2B). When we analysed the AURKA sequence using the bioinformatic algorithms, TargetP and MitoProt (Claros and Vincens, 1996; Emanuelsson et al., 2007), we identified a ‘cryptic’ targeting sequence from AA 7 – 55. That is, whereas the full length AURKA sequence was not predicted to localize to mitochondria, *in silico* removal of the first 6 AA from its N-terminus generated a predicted mitochondrial localization signal (probability in MitoProt, p>0.95, Figure 4B). To test the prediction that an N-terminally clipped version of AURKA, AURKAΔ6, should localize more strongly to the mitochondrial network than the full-length protein, we compared the localisation of AURKA-Venus and AURKAΔ6-Venus in fixed RPE-1 cells. Indeed, AURKAΔ6-Venus showed a more prominent co-localization with mitochondria, consistent with the presence of cryptic mitochondrial targeting information in the AURKA N-terminus (Figure 4C). Moreover, mitochondrial fragmentation was also enhanced in the presence of AURKAΔ6-Venus, supporting the idea that mitochondrial targeting of AURKA promotes mitochondrial fission (Figure 4D).

**Figure 4.**
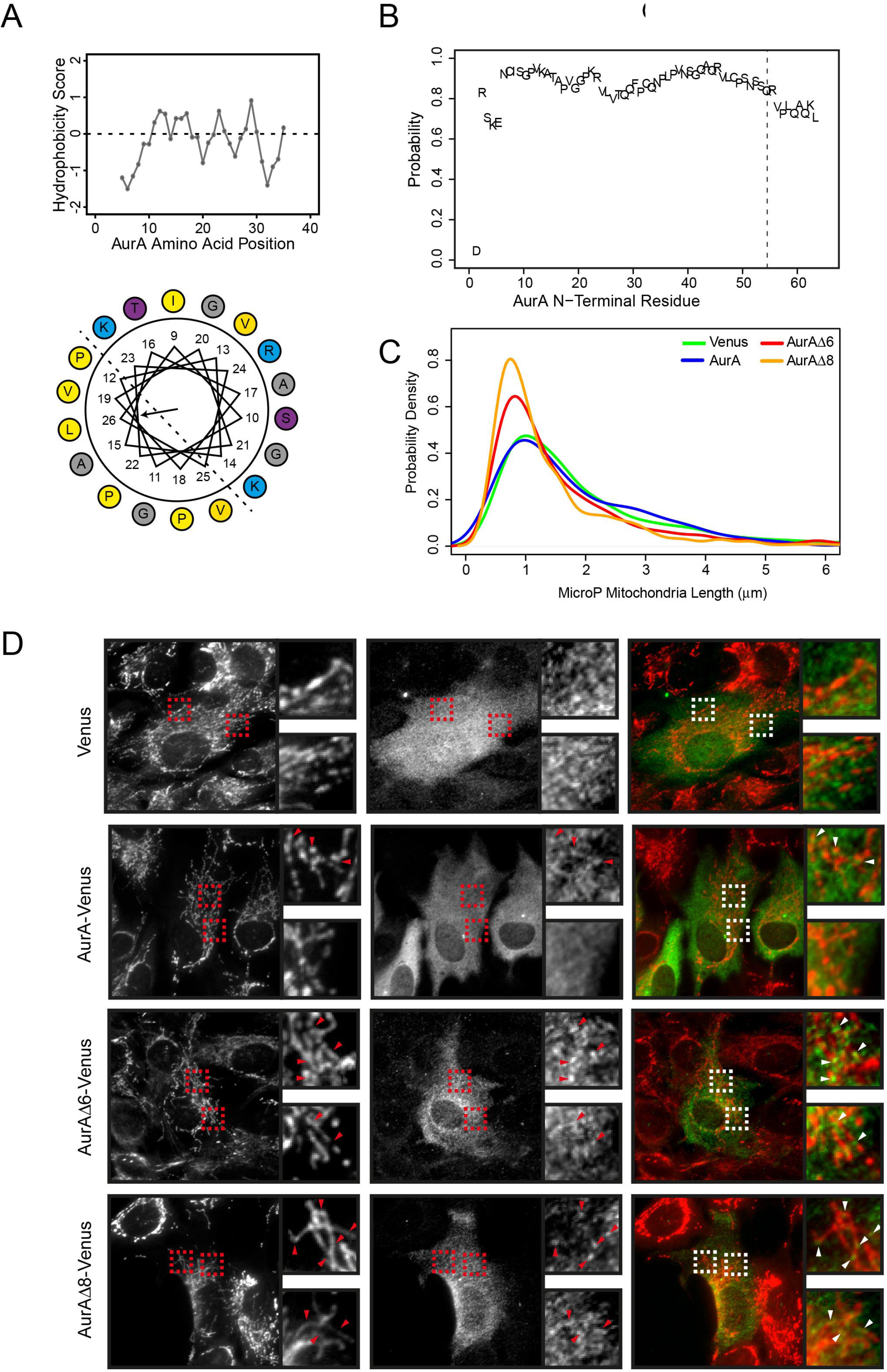
A cryptic mitochondrial targeting sequence resides in AURKA N-terminus. **A** Likely amphipathic helix in AURKA N-terminus revealed by Kyte-Doolittle plot (top panel) and Helical wheel projection (bottom panel) (Gautier et al., 2008). Yellow, hydrophobic; blue and purple, hydrophilic; grey, neutral. **B** MitoProt prediction of mitochondrial targeting probability (Claros and Vincens, 1996) for *in silico* sequential N-terminal truncations of AURKA (AurA). **C,D** RPE-1 cells were transfected with wild-type or N-terminally truncated versions of AURKA-Venus (Δ6, Δ8) and imaged 24h later after staining with MitoTracker™ for measurements of mitochondrial tubular length (**C**). Cells were then fixed and processed for IF with GFP and TOMM20 antibodies (**D**).

A large number of proteins are described to interact with AURKA (Shagisultanova et al., 2015). Some of these interactions may depend upon alternative conformations of the N-terminus. Thus, for example, the so-called A-box region can mediate interactions with either the APC/C co-factor Cdh1 or with Calmodulin, in a manner that may be regulated by phosphorylation on Ser51 (Crane et al., 2004; Plotnikova et al., 2010). We suggest that factors affecting the folding of the N-terminal region of AURKA could determine the localization and fate of the kinase by directing alternative interactors. One form of AURKA in this conformational space would display a functional MTS to mediate its mitochondrial targeting. This conformation would be favoured by AURKAΔ6.

Is AURKA on the surface of mitochondria, as predicted by its known role in promoting Drp1-mediated fission, or on the inside, as predicted by its MTS? Studies of mitochondrial regulation by mitotic cyclin-dependent kinase (Cdk) activity point to roles for cyclinB1-Cdk1 both at the mitochondrial surface, in regulating Drp1 activity (Kashatus et al., 2011), and – in a different study - inside the mitochondrial matrix, in regulating the respiratory chain via Complex I phosphorylation (Wang et al., 2014). The targeting sequence on AURKA suggests that it would be translocated into the mitochondria via the TOMM complex. Indeed, we have found direct interaction with the TOMM20 and TOMM70 members of this complex and also identified ATP5A subunits and other matrix components in our proteomic survey of AURKA interactors (Figure S1A).

The ‘MAGIC’ pathway recently described by Rong Li lab diverts aggregation-prone proteins into mitochondria by an unknown mechanism that results in their ubiquitin-independent clearance by mitochondrial proteases (Ruan et al., 2017). It is possible that AURKA is an aggregation-prone protein, given the unstructured nature of its extended N-terminus. Exogenously expressed protein would be more likely processed via this pathway than endogenous proteins, since it is more likely to be expressed in absence of the correct binding partners, and we certainly cannot exclude that this pathway is responsible for colocalization of AURKA with mitochondria. However, both the endogenous and exogenous AURKA is processed in this way and thereby contributes to the dynamics of the mitochondrial network.

Mitochondrial fragmentation is accompanied by increased glycolysis and decreased oxidative phosphorylation, metabolic changes that are thought to play a critical role in cell fate decisions (discussed in (Xu et al., 2013)). A switch to high glycolytic flux and low oxidative metabolism is a condition for reprogramming of iPS cells, with the state of high pluripotency being characterized by a fragmented mitochondrial network. It has been reported that AURKA is upregulated in reprogramming of iPS cells, although – curiously – inhibition of AURKA appears to enhance the process (Li and Rana, 2012). Our finding that the mitochondrial network is sensitive to AURKA levels in different cell types highlights the importance of elucidating the non-mitotic roles of AURKA in order to fully understand its contributions to cell proliferation and to cancer.

## Acknowledgements

Our thanks go to Helfrid Hochegger for sharing unpublished results, to David Nelson and Heike Laman for sharing protocols, to Jonathon Pines and Paola Marcos Casanova for help generating the endogenous AURKA-mVenus knock in, to Mingwei Min for help with data analysis and to Chris Connor and Begum Akman for practical assistance. This work was supported through funding to CL from Cancer Research UK (C3/A10239), Medical Research Council (MR/M01102X/1) and the Department of Genetics. DMG received support from Cancer Research UK (C3/A18795). RG was supported by a PhD studentship from the MRC and additional funding from the Department of Genetics, Cambridge Philosophical Society, Crane’s Charity and DRC. AMA is recipient of a Yousef Jameel Scholarship from the Cambridge International Trust. AB received an Erasmus placement (European Commission Lifelong Learning programme) and MPG was supported by Consolider (CSD2009-00016) and Ministerio de Ciencia e Innovación, Spain, through JdC and Jose Castillejo programs. JM is supported by the German Research Foundation (DFG) (Emmy Noether; MA 5831/1-1) and receives funding from the European Research Council (ERC) under the European Union’s Horizon 2020 research and innovation program (grant agreement no. 680042).

## Author contributions

RG carried out data acquisition and analysis and prepared figures. AMA and AB contributed specific experiments. MPG generated the RPE-1 FRT/TO AURKA-Venus line whilst JM generated endogenously tagged AURKA-mVenus. The project was conceived by CL and DMG and directed by CL. RG, JM, DMG, CL all contributed to writing and reviewing the manuscript.

